# Identification of novel enolase negative *Segatella copri* subspecies supports notion of *Segatella copri* speciation via alternative phosphoenolpyruvate synthesis pathways

**DOI:** 10.64898/2026.06.02.729653

**Authors:** Luke M. Bosnar, Steve Petrovski, Anya Shindler, Ashley E. Franks

## Abstract

**Background:** *Segatella copri* is characterised as a prominent glycolytic plant-based fiber utiliser within the human gut. My recent work has introduced a new interpretation of the positive impacts of plant-based polysaccharides on *S. copri*, as a significant negative relationship between *S. copri* and *Blautia* spp. was identified. The high rate of electron donor consumption by *Blautia* spp. indicated that competition for the electron donors, formate, ferredoxin and fumarate, could be the route of the negative relationship but would also explain the positive relationship with plant-based polysaccharides, as they are products of fiber fermentation intestinally.

**Methods:** Forty two genomes of *S. copri* were annotated via Prokka to identify alternative PEP pathways. Phylogenetics allowed effectively classification of the *S. copri* isolates into species and subspecies clusters. The sequence homology of nucleotides and proteins were analysed against the control, *S. copri* DSM 18205, to determine the level of conservation in the alternative phosphoenolpyruvate synthesis pathways.

**Results:** Enolase (*eno*) was not identified in the *S. copri* strains; JCM 13468, LKV-178-WT-2C, RHA03, RHA01 and RHA02, and the whole genome phylogenetic grouping of these strains, has proposed the existence of an *eno*(-) subspecies of *S. copri*. This work furthered this idea by identifying alternative PEP pathways from formate, ferredoxin and fumarate, which were the most conserved in the *eno*(-) *S. copri* genomes.

**Conclusion:** This work has provided rationale to why enolase may not be present within the eno(-) *S. copri* isolates and have shown that these alternative PEP synthesis pathways could negate the requirement of enolase in cells and may be factor in evolution of *S. copri* metabolism.

## Introduction

The phosphoenolpyruvate-pyruvate-oxaloacetate (PPO) node consists of the pyruvate-phosphoenolpyruvate (Pyr-PEP) and oxaloacetate-phosphoenolpyruvate (OAA-PEP) substrate cycles, also known as futile cycles, and represents a dynamic interplay between metabolic pathways in which pyruvate and oxaloacetate are recycled to generate phosphoenolpyruvate (PEP) (Sharma et al. 2024). Futile cycles typically yield no net gain of products as the adenosine triphosphate (ATP) produced is often utilised during the cycle (Sharma et al. 2024; Brownstein et al. 2022). Whilst substrate cycles can be metabolic dead ends, certain pathways have been observed to utilise alternative phosphate donors in leu of ATP (Qian et al. 2006; Auger et al. 2021). The presence of free phosphates has been found to affect the utilisation of guanosine triphosphate (GTP) or inorganic phosphates (Pi) during the biosynthesis of PEP and may result in a net ATP gain from both the Pyr-PEP and OAA-PEP substrate cycles (Qian et al. 2006; Auger et al. 2021; Koendjbiharie et al. 2021; Cline et al. 2011).

The Pyr-PEP substrate cycle is a critical metabolic junction that involves the recycling of pyruvate to produce PEP, commonly via reutilising the pyruvate produced during the dephosphorylation of PEP by pyruvate kinase (*pyk*) (Figure 1) (Koendjbiharie et al. 2021; Cline et al. 2011; Llamas-Ramirez et al. 2020; Koendjbiharie et al. 2018). While the OAA-PEP substrate cycle requires the action of *pckA* to produce PEP from oxaloacetate but does not acquire oxaloacetate during the dephosphorylation of PEP by *pyk* (Figure 1) (Koendjbiharie et al. 2021; Llamas-Ramirez et al. 2020). Instead, the completion of OAA-PEP substrate cycle occurs via conversion of pyruvate to malate by the malic enzyme (*maeB*), and the conversion of malate to oxaloacetate by malate dehydrogenase (*mdh*) (Figure 1) (Koendjbiharie et al. 2021).

**Figure 1:**
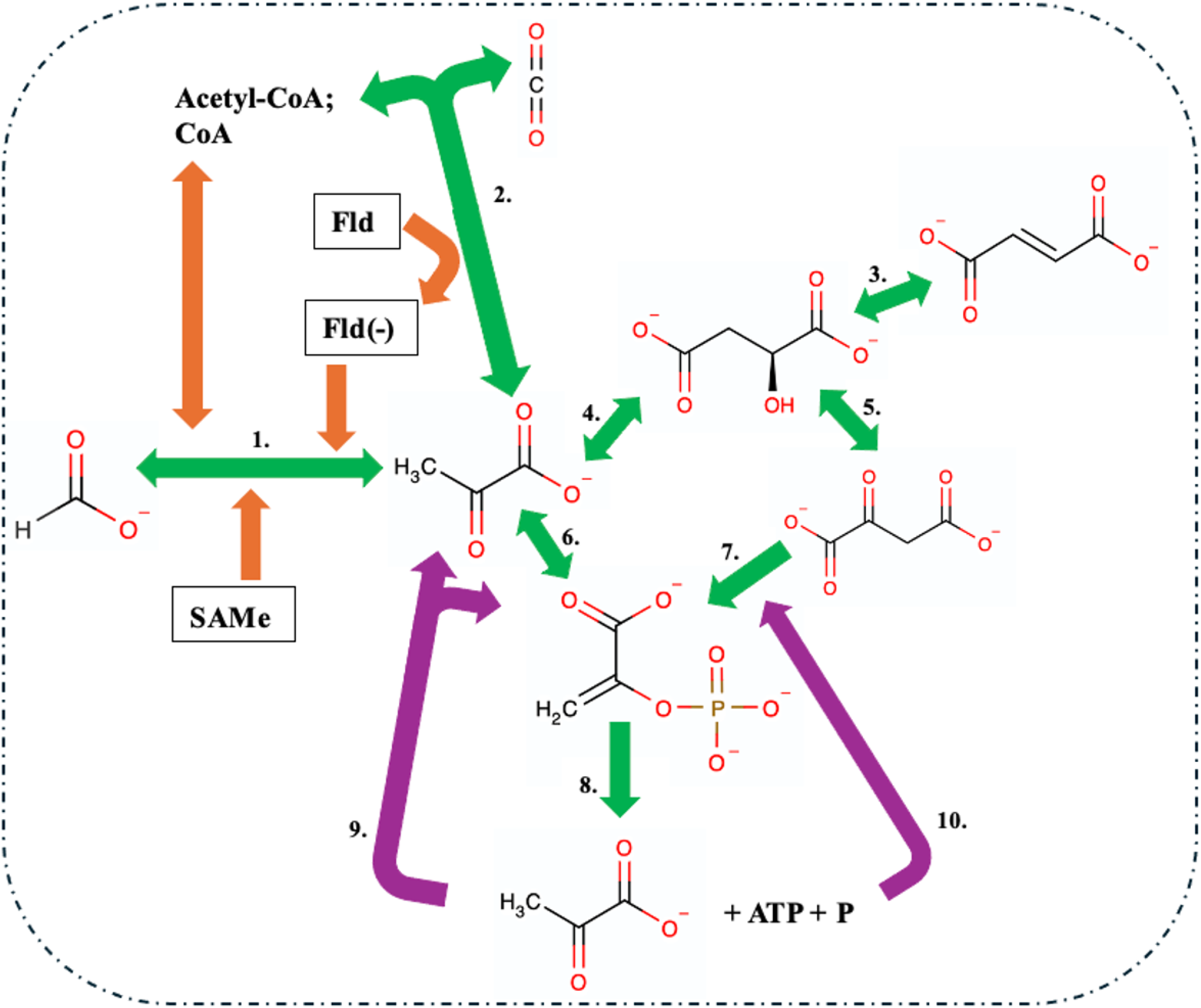
The Pyruvate-Oxaloacetate-Phosphoenolpyruvate (PPO) node could be accessed by the production of pyruvate or oxaloacetate via pyruvate, phosphate dikinase (*ppdK*) and phosphoenolpyruvate carboxykinase (*pckA*), respectively. Within the DSMZ type-strain *Segatella copri* DSM 18205, genes encoding enzymes with the capability of accessing pyruvate or oxaloacetate from the utilisation of formate, ferredoxin and fumarate were identified, as well as *ppdK* and *pckA*. **Legend:** 1. Formate acetyltransferase (*pflB*); 2. Pyruvate:ferredoxin oxidoreductase (PFOR); 3. Fumarate hydratase (*fumA*); 4. Malic enzyme (*maeB*); 5. Malate dehydrogenase (*mdh*); 6. Pyruvate, phosphate dikinase (*ppdK*); 7. Phosphoenolpyruvate carboxykinase (*pckA*); 8. Pyruvate kinase (*pyk*); 10. The shuffling of pyruvate back to *pflB* and *ppdK;* 11. The shuffling of adenosine triphosphate (ATP) and a free phosphate (P) back to *pckA*.

The PPO node can be accessed by multiple metabolites but occurs only via the conversion to pyruvate or oxaloacetate first, by *ppdK* and *pckA,* respectively (Figure 1) (Koendjbiharie et al. 2021; Cline et al. 2011; Llamas-Ramirez et al. 2020). Pyruvate and oxaloacetate are present in cells as intermediate products of metabolism and serve as substrates in various metabolic processes, including glycolysis, gluconeogenesis, tricarboxylic acid cycle, Pyr-PEP and OAA-PEP cycles (Yuan et al. 2022; Maleki et al. 2017). Due to the high demand by a number of key metabolic pathways for pyruvate and oxaloacetate, intracellular levels are tightly regulated by the conversion of other molecules, such as lactate or malate, respectively, to maintain cellular pyruvate or oxaloacetate levels (Yuan et al. 2022; Maleki et al. 2017; Kang et al. 2024). Moreover, pyruvate and oxaloacetate can be interchangeably converted via malate to regulate intracellular levels and could play a role in controlling substrate levels within the PPO node (Figure 1) (Koendjbiharie et al. 2021; Yuan et al. 2022).

Microbes are capable of utilising either formate, fumarate or ferredoxin to produce pyruvate independently, with the capacity to produce oxaloacetate from fumarate via the conversion to malate first (Figure 1) (Luo et al. 2023; Bonitatibus et al. 2025; Wei et al. 2021; Asanuma et al. 1999). Formate can be converted directly to pyruvate by the pyruvate formate-lyase enzyme complex (*pflAB*), but the activation of *pflB* requires the cofactors, S-adenosylmethionine (SAMe) and a reduced flavodoxin (Figure 1) (Zhang et al. 2024; Crain and Broderick 2014; Broderick et al. 2000; Cáceres et al. 2023). Various microbes do have the capacity to synthesise SAMe via S-adenosylmethionine (*metK*) and flavodoxin via (*fldA*) and these genes may have elevated transcription during *pflB* reactions (Figure 1) (Troitzsch et al. 2024; Boswinkle et al. 2022). The reduction of flavodoxin is also necessary for *pflB* activation and can occur through the function of pyruvate:ferredoxin oxidoreductase (PFOR) (Figure 1) (Bonitatibus et al. 2025; Zhang et al. 2024). However, the reduction of flavodoxin via PFOR is a thiamine pyrophosphate (TPP) dependent reaction and suggests that a microbe reducing flavodoxin via PFOR to activate *pflB* would require TPP (Bonitatibus et al. 2025; Zhang et al. 2024; Katsyv et al. 2021).

Moreover, fumarate cannot be directly converted to pyruvate and first must be converted to malate by fumarate hydratase (*fumA*) and then pyruvate by *maeB* (Figure 1) (Luo et. al 2023; Bonitatibus et al. 2025; Wei et al. 2021). Alternatively, the malate produced by *fumA* and *maeB* from fumarate and pyruvate, respectively, could be converted to oxaloacetate instead by *mdh*, and these reactions may be dependent upon pyruvate and oxaloacetate levels (Figure 1) (Koendjbiharie et al. 2021; Yuan et al. 2022; Luo et al. 2023; Bonitatibus et al. 2025; Wei et al. 2021). While ferredoxin is not converted but instead utilised as an electron donor in the production of pyruvate from CO_2_ and acetyl-CoA by PFOR (Bonitatibus et al. 2025). The utilisation of ferredoxin by PFOR has predominantly been reported in the pyruvate catabolic direction, but the capacity for this pathway to operate bidirectionally has been seen in autotrophic microbes (Bonitatibus et al. 2025; Furdui and Ragsdale 2000).

PEP, pyruvate and oxaloacetate are intermediates in multiple metabolic pathways, including glycolysis, gluconeogenesis and the TCA cycle, with enolase (*eno*) being a key enzyme linking these processes (Koendjbiharie et al. 2021). The PPO node is capable of accepting PEP, pyruvate or oxaloacetate from these metabolic pathways, whilst simultaneously generating PEP and the PEP derivatives, pyruvate, ATP and Pi, to be utilised by or activate other metabolic pathways (Koendjbiharie et al. 2021; Cline et al. 2011; Llamas-Ramirez et al. 2020; Koendjbiharie et al. 2018; Saur et al. 2005). Gluconeogenesis can be activated by the PPO node to facilitate glucose production, with PEP being converted by *eno* to 2-phosphoglycerate, consuming Pi but producing a pyruvate molecule, that could be reutilised in the PPO node (Koendjbiharie et al. 2021; Gupta et al. 2021; Melkonian et al. 2019). Additionally, acitivity of the glycolytic and the TCA cycles can provide substrate for the PPO node, via the pyruvate and Pi released from PEP during glycolysis and Pi during the conversion of PEP to oxaloacetate in the TCA cycle (Stark et al. 2009; Chaudhry et al. 2018; Haddad et al. 2023). The ability to recycle substrates from other metabolic pathways to regenerate PEP confers a cellular benefit due to activation of various metabolic pathways as well as an additional energy source and phosphate donor (Stark et al. 2009; Hu et al. 2025; Delbaere et al. 2004).

The production of PEP without enolase is known to occur by three microbial enzymes, phosphoenolpyruvate synthase (*ppsA*), phosphoenolpyruvate carboxykinase (*pckA*) and pyruvate, phosphate dikinase (*ppdK*) (Koendjbiharie et al. 2021; Long et al. 2017). The upregulation of gene transcription has been associated within nutrient limited intestinal environments and reported to have negative health outcomes, with the elevated transcription of *ppsA, pckA* and *ppdK* associated with contributing to the survival of pathogens (Bertin et al. 2014; Rohde et al. 2012; Basu et al. 2018). Alternatively, the transcription of *ppdK* has been associated with microbial symbiosis amongst commensal bacteria via a series of *in vitro* co-culturing of *ppdK* containing and *ppdK*(-) mutants of *Acetobacter fabarum*, with a wild-type *Levilactobacillus brevis* (Sommer and Newell 2019). The *in-vitro* cross-feeding between *A. fabarum* and *L. brevis* revealed that *ppdK* allowed the lactate produced by *L. brevis* to be converted to pyruvate, which activates gluconeogenesis in *ppdK*(+) *A. fabarum* via the subsequent conversion of pyruvate to PEP via *ppdK* (Sommer and Newell 2019). The metabolic cooperation benefit may not be limited to microbes, as *ppdK* is consistently upregulated in nutrient limited environments, and may serve as a route to PEP and glucose production from bacterial metabolites and or non-carbohydrate sources, alleviating intestinal PEP/glucose levels regardless of intestinal nutrient content (Rowland et al. 2018; Sommer and Newell 2019).

In saccharolytic bacteria, surplus PEP may enhance the transport of sugars into the cell, via the phosphotransferase system (PTS), promoting efficient transport and utilisation of monosaccharides (Delbaere et al. 2004; Siebold et al. 2001). Conversely, bacteria lacking a complete PTS apparatus may benefit from the increased activation of PEP dependent metabolic pathways induced by the increased levels of PEP (Hu et al. 2025; Lim et al. 2007; Minges et al. 2017).

Although pyruvate and oxaloacetate levels in the intestines have been studied, a link between formate, fumarate or ferredoxin and the PPO node within the intestinal microbiome has not been suggested, despite it being possible to acquire pyruvate or oxaloacetate from these compounds through known enzymes (Hughes et al. 2017; Ternes et al. 2022; Papila et al. 2024; Garg et al. 2018; Koendjbiharie et al. 2021; Koendjbiharie et al. 2018). Formate, fumarate and ferredoxin are mainly associated with acetogens and methanogens, which utilise them for the Wood-Ljungdahl pathway (WLP) or methanogenesis, respectively (Luo et al. 2023; Bonitatibus et al. 2025; Wei et al. 2021; Asanuma et al. 1999). No other direct link with formate or ferredoxin and other microbes remains unreported, while fumarate is involved in the anaerobic respiration of some bacteria (Bui et al. 2019; Laverde Gomez et al. 2019; Kelly et al. 2022; Karekar et al. 2022).

In an environment that supports acetogens and methanogens such as the gastrointestinal tract, electron carrying molecules, such as formate, fumarate and ferredoxin are commonly consumed and accompanied by increased methane levels, directly by methanogens and indirectly by acetogens (Bonitatibus et al. 2025; Wei et al. 2021; Kumpitsch et al. 2021; Triantafyllou et al. 2014; Suri et al. 2018; Mutuyemungu et al. 2023). While acetogens are often linked to reduced methane production, the acetate produced via acetogenesis can be utilised by methanogens and can positively impact methanogenesis (Kumpitsch et al. 2021; Sun et al. 2021; Karakashev et al. 2006; Prochazkova et al. 2023; Wang et al. 2023). Methanogens also use hydrogen as an electron donor so acetogens may not always reduce methanogenesis when formate, fumarate or ferredoxin and hydrogen are present (Kumpitsch et al. 2021; Sun et al. 2021; Karakashev et al. 2006; Prochazkova et al. 2023; Wang et al. 2023).

The levels of acetate production via acetogens in response to formate utilisation can be significantly higher compared to CO_2_ and H_2_ fuelled acetogenesis due to formate-acetogenesis being more energy efficient and having a quicker reaction rate than CO_2_ and H_2_ (Moon et al. 2021; Kerby and Zeikus 1987). Moreover, the products of formate-acetogenesis are often CO_2_ and H_2_, which indicates that formate-acetogenesis may enhance cross-feeding with other acetogens or methanogens (Moon et al. 2021; Kerby and Zeikus 1987). The potential role of formate-acetogenesis and acetogen-methanogen cross-feeding suggests that a microbe which can consume formate without producing greater levels of acetate CO_2_ and H_2_ than formate-acetogenesis could disrupt acetogen-acetogen and acetogen-methanogen cross-feeding events fuelled via formate (Moon et al. 2021; Kerby and Zeikus 1987).

A number of studies have reported an inverse relationship with the genus’, *Segatella* spp., *Fibrobacter* spp. and *Ruminococcus* spp., and methanogenesis in dairy cows, suggesting that species within these genus’ may be negatively correlated with methane levels due to competition for electron carrying molecules (Kumpitsch et al. 2021; Sun et al. 2021; Karakashev et al. 2006; Prochazkova et al. 2023; Aryee et al. 2023). The negative correlation between *Segatella* spp. and methane production was also observed in Colombian buffalo, where increased abundance of *Segatella* spp. corresponded to lower methane levels, potentially reflecting the association of a diminished methanogenic activity with an increased relative abundance of *Segatella* spp. (Kumpitsch et al. 2021; Sun et al. 2021; Karakashev et al. 2006; Prochazkova et al. 2023; Aryee et al. 2023; Aguillar et al. 2020). Although methane production does not directly indicate the utilisation of either formate, fumarate or ferredoxin, past research has examined the interactions between acetogens and methanogens with formate and ferredoxin through growth kinetic analysis (Furdui and Ragsdale 2000; Dietrich and Muller 2023; Trischler et al. 2022).

Formate supplemented cultures demonstrated that *Blautia* spp., *B. hydrogenotrophica, B. luti* and *B. wexlerae*, preferentially utilised formate over hydrogen as an electron donor during the WLP *in vitro* (Trischler et al. 2022). The strength of the association between the acetogenic *Blautia* spp. and formate was further highlighted during a co-culturing experiment where the formate produced by *R. bromii* was completely consumed by *B. hydrogenotrophica* by the end of the co-culturing experiment (Laverde Gomez et al. 2019). The *in vitro* co-culturing analysis did not include ferredoxin or fumarate utilisation by the *Blautia* spp. but did show the increased expression of *fumA* and PFOR within the acetogenic co-cultures (Laverde Gomez et al. 2019). Moreover, formate utilisation by the methanogenic bacteria, *Methanobrevibacter smithii,* has also been observed to be elevated in the presence of acetate, suggesting that formate competition may be increased by the relative abundance of both acetogenic and methanogenic bacteria (Wang et al. 2023).

While the effect of intestinal formate, ferredoxin or fumarate on *Segatella* spp. has not been explored, research has shown formate can enhance growth in an *in vitro* simulated infant gut model (Parkar et al. 2021). Significant rises in the relative abundance of *Segatella* spp. in the ‘Digesta Control’ groups were observed, with the control media containing higher formate levels than the test media (Parkar et al. 2021). If a microbe possesses *pflAB* to convert formate to pyruvate and then PEP via *ppdK*, the elevated formate levels may enable entry into a PPO substrate cycle via pyruvate and potentially account for the significant rises in the relative abundance of *Segatella* spp. in the ‘Digesta Control’ groups (Koendjbiharie et al. 2021; Cline et al. 2011; Llamas-Ramirez et al. 2020; Koendjbiharie et al. 2018; Parkar et al. 2021). The capacity to utilise formate to produce PEP would provide increased microbial fitness via increased metabolic activity, energy and phosphate donation, yet would also leave the microbe susceptible to heavy formate competition within the intestines (Stark et al. 2009; Hu et al. 2025; Delbaere et al. 2004).

While recent research has looked into the capacity of *Segatella copri* to produce formate and fumarate during *in vitro* growth on glucose, this work did not identify if the formate or fumarate produced was also being utilised by *S. copri* (Franke and Deppenmeier 2018; Huang et al. 2021). As this research did not set out to investigate the metabolism of *S. copri* associated with formate or fumarate, the levels of pyruvate were not recorded and therefore the rate at which formate, or fumarate is being produced cannot be aligned with the rate at which these compounds are converted to pyruvate (Huang et al. 2021). However, during the Franke and Deppenmeier study of *S. copri* DSM 18205 it was identified that both pyruvate and PEP were the two key central intermediates during the *in vitro* growth experiment and transcriptomic analysis showed that *pckA* was one of the most transcribed genes during the log phase (Franke and Deppenmeier 2018).

*Segatella* spp. metabolism via the PPO node can be theorised through the homeostatic role of the PPO node in insulin sensitivity within the pancreas, and the host related health outcomes associated with PPO metabolism (Stark et al. 2009; Merrins et al. 2022; Jessinkey et al. 2019; Xiao et al. 2024; Zang et al. 2024; Abdelsalam et al. 2023). *S. copri* abundance has been observed to be associated with fluctuating insulin sensitivity but also improved glucose metabolism in two separate diabetic animal models, with the regulatory mechanism of *S. copri* abundance suggested to be the increase of glucose homeostatic factors in the intestines including succinate, the secretion of glucagon-like peptide-1 (GLP-1) and upregulation of Farnesoid X receptor (FXR) (Yang et al. 2024; Pean et al. 2020).

If present in a relative abundance intestinally, a microbe utilising PPO node as a central metabolism could impact glucose homeostasis by reducing glucose competition via PEP mediated gluconeogenesis and in turn limiting the activation of the glucose regulation cascade (Stark et al 2009; Merrins et al. 2022; Jessinkey et al. 2019; Xiao et al. 2024; Zang et al. 2024; Abdelsalam et al. 2023; Yang et al. 2024; Pean et al. 2020). *S. copri’s* activation of the PPO node may result in the increased export of PEP into the intestinal environment and allow gluconeogenic bacteria to facilitate glucose synthesis (Koendjbiharie et al. 2021; Melkonian et al. 2019). Hypothetically, the access to and exportation of PEP by the PPO node by a relatively abundant intestinal microbe within the intestines, could facilitate a community wide PEP mediate gluconeogenesis, which would significantly reduce glucose competition intestinally (Koendjbiharie et al. 2021; Melkonian et al. 2019).

The relationship between *S. copri* and glucose regulation is characterised as dynamic, with the relative abundance of *S. copri* associated with both high and low blood glucose (Franke and Deppenmeier 2018; Kovatcheva-Datchary et al. 2015; Liu et al. 2023). The varied impact of *S. copri* colonisation within the intestine on glucose metabolism may reflect environmental factors, potentially influenced by the catalytic activity of *S. copri* within that intestinal setting (Xiao et al. 2024; Kovatcheva-Datchary et al. 2015; Golisch et al. 2024). Prior research proposed succinate production by *S. copri* as a possible mechanism for the regulation of glucose metabolism in diabetic mice, and the production of succinate is directly impacted by the flow of metabolic intermediates from the PPO node (Koendjbiharie et al. 2021; Koendjbiharie et al. 2018; Yang et al. 2024; De Vadder et al. 2016).

Furthermore, *S. copri* is recognised as a key intestinal microbe involved in human health and has been linked to various diseases, including diabetes, arthritis and allergies, although the role *S. copri* plays within these disorders is unknown (Xiao et al. 2024; Zang et al. 2024; Abdelsalam et al. 2023). A potential link between *S. copri* and diabetes may be relevant to the glucose regulatory function of the PPO node (Koendjbiharie et al. 2021; Koendjbiharie et al. 2018; Yang et al. 2024; De Vadder et al. 2016). The hypothesis that *S. copri* is producing elevated levels of PEP could also explain the complex involvement of *S. copri* in inflammatory intestinal diseases, as PEP has been identified to contribute to intestinal inflammation by interfering with Ca^2+^ signalling, whilst also reducing viral infections and intestinal inflammation (Soto-Heredero et al. 2020; Moreno-Felici et al. 2019; Zhao et al. 2023). Alternatively, *S. copri* access to the PPO node could result in reduced levels of intestinal PEP if PEP and PEP intermediates are reutilised within the PPO node rather than being exported (Koendjbiharie et al. 2021; Koendjbiharie et al. 2018). *S. copri’s* ability to obtain pyruvate and oxaloacetate from formate, fumarate or ferredoxin utilisation whilst accessing the PPO node could lead to increased PEP synthesis by *S. copri,* which may impact levels of PEP and PEP derivatives within the intestine (Koendjbiharie et al. 2021; Koendjbiharie et al. 2018; Montal et al. 2015; Oliphant et al. 2019).

Whilst PEP’s role in inflammatory disease varies, multiple beneficial associations between PEP and intestinal epithelia have been reported, including T-cell function and maintenance of gut barrier integrity (Ho et al. 2015; Yu et al. 2023; Yin et al. 2024). Although T-cell regulation and intestinal epithelial maintenance have been correlated with the relative abundance of *S. copri*, the production and export of PEP or PEP derivatives by *S. copri* within the intestines has not been investigated as a mechanism in the context of intestinal inflammatory disease (Xiao et al. 2024; Zang et al. 2024; Abdelsalam et al. 2023; Bhutta et al. 2024). The increased synthesis of PEP and PEP intermediates by *S. copri* intestinally may impact the host cells by modulating the intestinal microbiome but may also lead to the over proliferation of microbes accessing exported PEP (Koendjbiharie et al. 2021; Koendjbiharie et al. 2018; Bhutta et al. 2024; Den Besten et al. 2013).

Despite *S. copri* being linked to various disorders, its exact function in the gut and host remains unclear. Studies have shown that the increased relative abundance of *S. copri* and PEP correlate to increased intestinal permeability (Yin et al. 2024; Zhang et al. 2025). Although increased intestinal colonisation of *S. copri* is associated with increased intestinal permeability, the action of *S. copri* upon intestinal permeability has not been established (Nieman et al. 2025; Lo Presti et al. 2023; Chen et al. 2021). Elevated intestinal PEP has been clearly connected to increased activity of epithelial phosphoenolpyruvate kinase (PEPCK), which promotes both the conversion between PEP and oxaloacetate, and heightened permeability and inflammation (Yin et al. 2024). Given the link between PEPCK transcription, and PEP and oxaloacetate, PEPCK may serve as a biomarker for intestinal PEP and oxaloacetate levels, potentially offering insights into microbial and epithelial interactions mediated by PEP and oxaloacetate (Yin et al. 2024). The production and export of PEP intestinally by bacteria is challenging to measure due to the high demand and rapid conversion potential of PEP and the PEP intermediates within the intestinal environment, as additionally various cell recycling mechanisms exist that enable the reutilisation of PEP and PEP derivatives to regenerate PEP (Koendjbiharie et al. 2021; Koendjbiharie et al. 2018; Montal et al. 2015; Sauer et al. 2005).

Multiple studies have examined the specific roles of *S. copri* in the gut and indicated that substrate levels within the intestines may influence energy metabolism in *S. copri*, rather than solely genetic differences; for example, increased formate, fumarate or ferredoxin concentrations may provide alternative metabolic pathways (Ternes et al. 2022; Xiao et al. 2024; Zang et al. 2024; Abdelsalam et al. 2023). In order to investigate the genomic possibility of the PPO node being accessed via formate, fumarate or ferredoxin utilisation within *S. copri*, the genes encoding the enzymes for formate fumarate and ferredoxin utilisation, and the subsequent conversions of formate, fumarate and ferredoxin to pyruvate and oxaloacetate, pyruvate and oxaloacetate to PEP, and PEP dephosphorylation to ATP and pyruvate were annotated and assessed within *S. copri* genomes accessed via NCBI genome database (https://www.ncbi.nlm.nih.gov) (Koendjbiharie et al. 2021; Koendjbiharie et al. 2018; Montal et al. 2015; Sauer et al. 2005).

## Materials and Methods

### *Segatella copri* DSM 18205 genomic analysis

The NCBI reference strain, *S. copri* DSM 18205, which has been previously utilised during *in vitro* growth experiments, was downloaded from the NCBI genome database (https://www.ncbi.nlm.nih.gov/biosample/SAMN20222705/) (Franke and Deppenmeier 2018; Huang et al. 2021). In order to examine the *S. copri* DSM 18205 genome for the presence of enzymes and metabolic pathways associated with PEP synthesis from sources alternative to glucose, the assembled genome (.fastq format) was analysed via the gapseq metabolic identifying pipeline (Zimmermann et al. 2021). The *S. copri* DSM 18205 genome was then run through the prokaryotic genome annotating pipeline, Prokka, and was identified to lack a functional and annotated *eno* (Seemann 2014). The investigation into the metabolic capacity of *S. copri* DSM 18205 included alternative genes involved in PEP synthesis (Seemann 2014).

### *Segatella copri* isolate selection criteria

The selection process for isolates included in this study, involved the recent taxonomic classification of the isolate as a member of the *S. copri* species. The location and date of isolation, and other strain information including isolate assembly accession numbers are available in Supplementary Sheet 1, and all isolates were accessed from the NCBI genome database.

### Whole-genome based phylogenetic tree of *Segatella copri* isolates

The forty one *S. copri* isolate query genomes were accessed from the NCBI genome database to be aligned with the NCBI reference *S. copri* DSM 18205; JCM 13468, N-01, BVe41219, CLA-KB-H127, MSK.21.28, YF2, iAP1411, MCC688, Ind63-31, AM22-1, AF12-50, RPG01, RA-N001-13, SG-1969, H-2477, F3-7, HDE06, Indica, Y7XP, BgG5_1, LKV-178-WT-2C, RHA01, RHA02, RHA03, N115-20, RPB01, HDC01, RPA01, HDD04, RPG01, RPF01, RPC01, Ind117-13, Ind117-18, Ind63-40, Ind63-29, N016-9, Ind117-32, Ind63-37, HDD05, F3-75 and HDE04 (https://www.ncbi.nlm.nih.gov/home/genomes/). The isolate *S. copri* JCM 13468 is a duplicate strain of *S. copri* DSM 18205 and was utilised as a positive control in this analysis. To determine the relatedness of each *S. copri* isolate, as well as the genetic similarity of each isolate to the reference *S. copri* DSM 18205, a whole genome phylogenetic tree was produced via the Type (Strain) Genome Server (https://tygs.dsmz.de).

### Identifying conserved alternate phosphoenolpyruvate synthesis pathways across divergent *Segatella copri* isolates

The sequences pertaining to the genes involved in the alternative PEP synthesis from formate, fumarate, ferredoxin, pyruvate and oxaloacetate were extracted from the nucleotide and protein FASTA files with annotated sequences (*.fnn* and *.faa*, respectively) produced by Prokka. The respective sequences of the genes of the query and reference genome were aligned via the NCBI sequence blastn and blastp pipelines, which produced a percentage identity (PI) for sequence homology (https://blast.ncbi.nlm.nih.gov/BlastAlign.cgi).

### Data availability and supplementary material statement

All relevant sequence data utilised in this study is available via the NCBI accession numbers present in the supplementary material (Supplementary Sheet 1).

## Results and Discussion

### Identification of incomplete glycolytic, gluconeogenic and tricarboxylic acid pathways in *Segatella copri* DSM 18205

Despite having various enolase enzyme and I-V glycolysis pathway queries during the gapseq analysis of *S. copri* DSM 18205, no *eno* was observed within the *S. copri* DSM 18205 genome during the analysis of the Prokka annotation report (Table 1). Due to the importance of *eno* for cells to complete glycolysis and gluconeogenesis, the lack of *eno* indicates the likely incapacity of *S. copri* DSM 18205 to complete glycolysis or gluconeogenesis (Table 1) (Koendjbiharie et al. 2021). Described as a prominent fiber utiliser, *S. copri* DSM 18205 not containing an *eno* will require further investigation to determine if this trait is present in other *S. copri* genomes (Koendjbiharie et al. 2021; Blanco-Miguez et al. 2023).

**Table 1:**
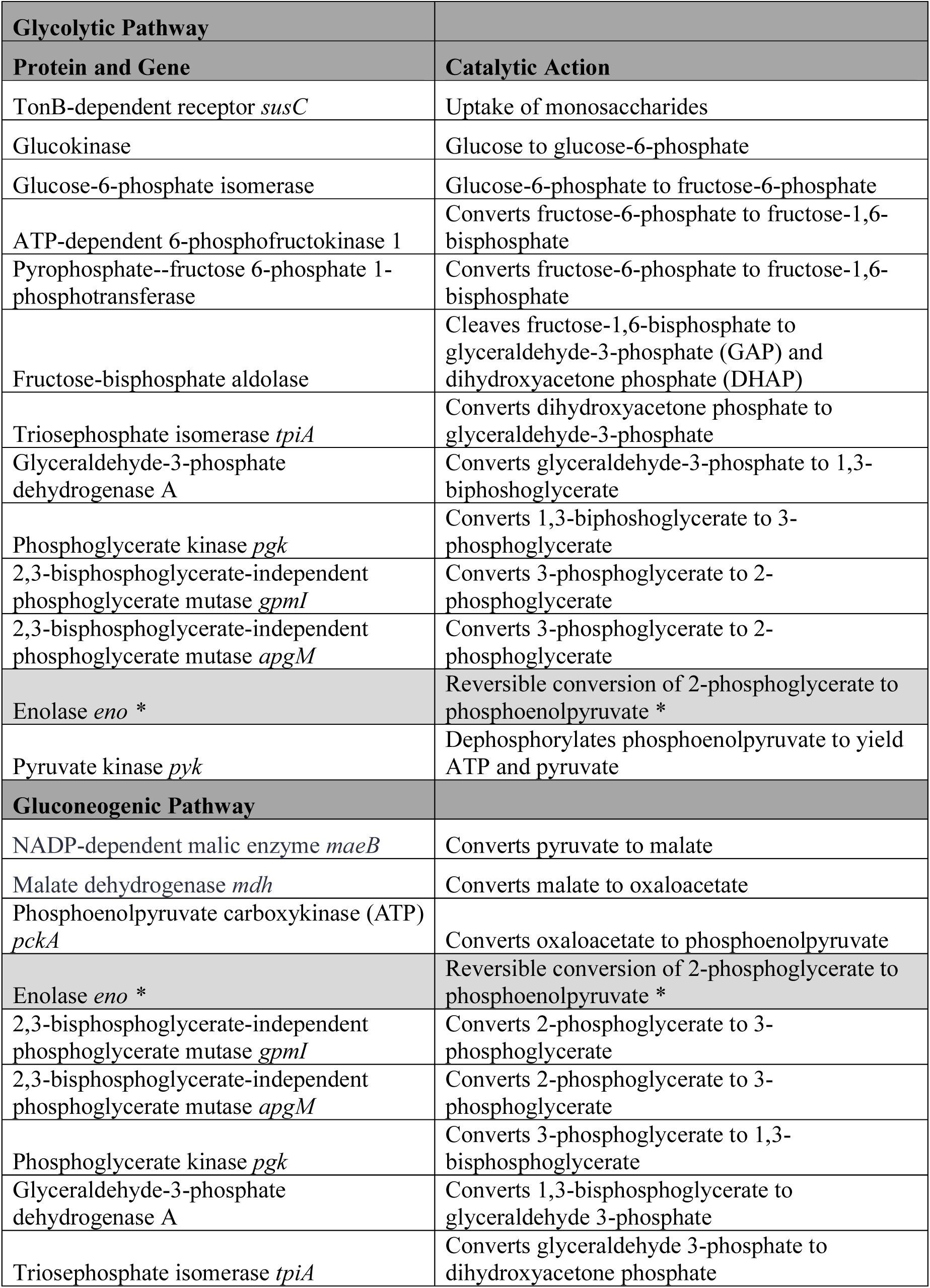
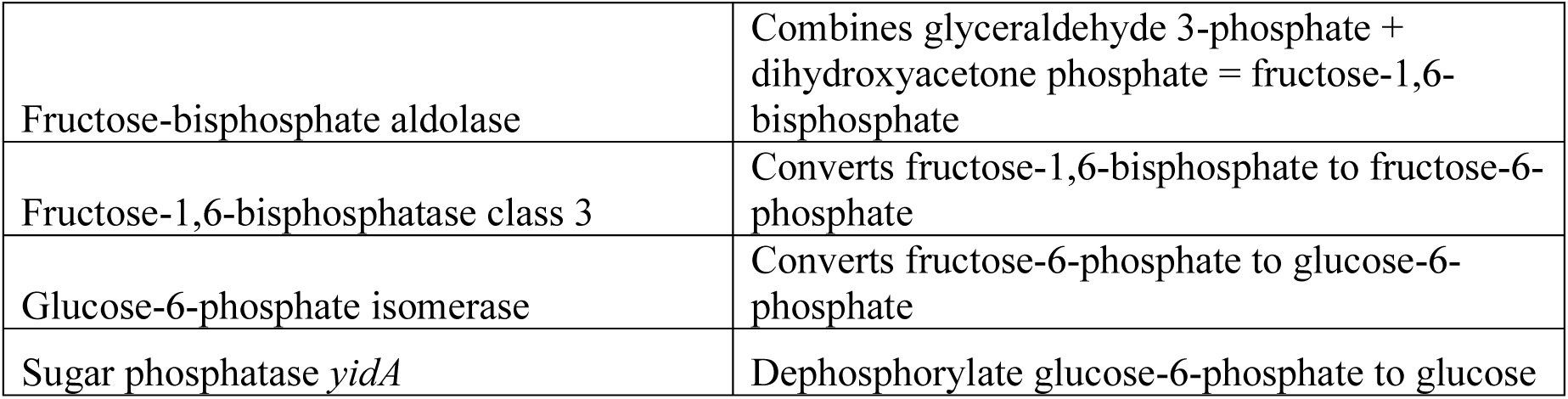
Through the analysis of the Prokka genome annotation of the DSMZ type-strain, *Segatella copri* DSM 18205, an enolase (*eno*) was not identified, despite the remaining genes involved in the glycolytic and gluconeogenic pathways being present. Absent genes and catalytic function were shaded with a *

Further investigation into the metabolic potential of *S. copri* DSM 18205, identified that the TCA pathway was not present in its entirety, as genes encoding the enzymes alpha-ketoglutarate dehydrogenase, succinyl-CoA synthetase and succinate dehydrogenase, were not present within the *S. copri* DSM 18205 genome (Table 2) (Koendjbiharie et al. 2021). Previously described as having the capacity to produce succinate, the lack of succinyl-CoA synthetase within the *S. copri* DSM 18205 genome, may not only suggest an inability to complete the TCA cycle but also that an alternate route to succinate production could exist in *S. copri* DSM 18205 (Table 2) (Koendjbiharie et al. 2021; Yang et al. 2024).

**Table 2:**
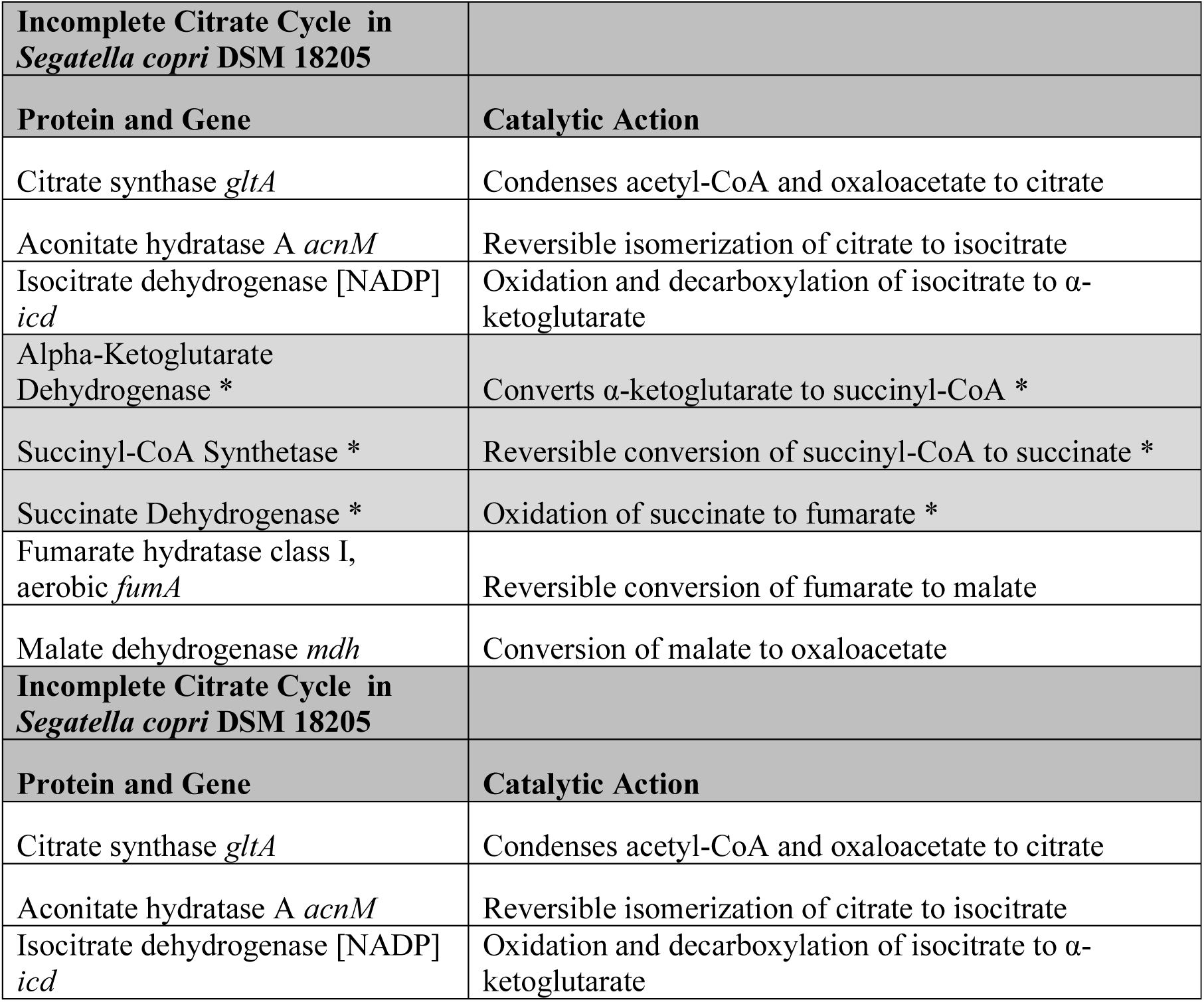
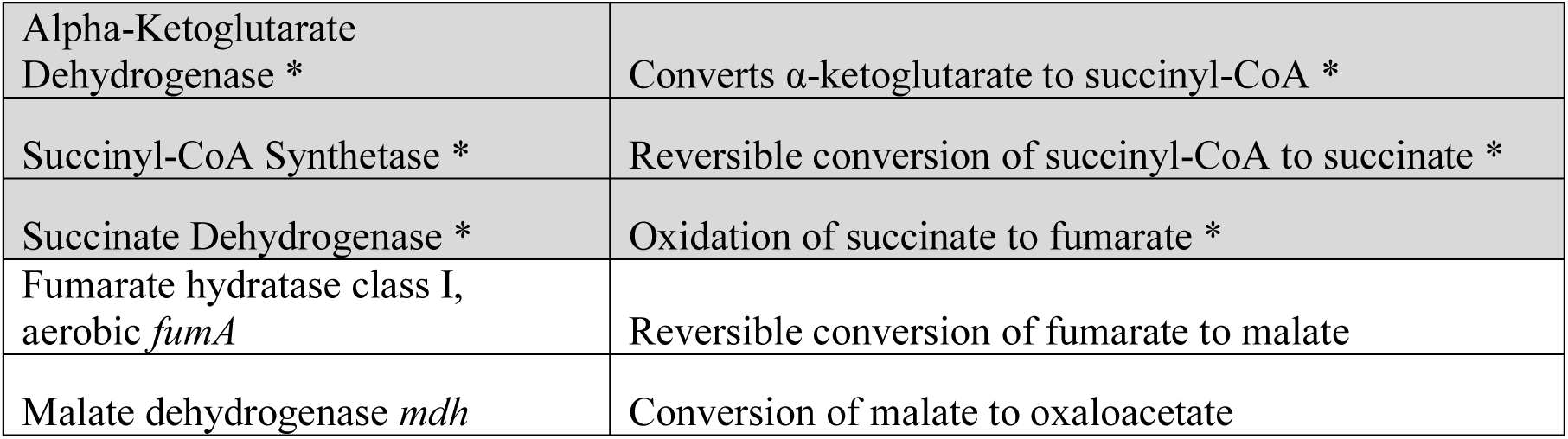
Through the analysis of the Prokka genome annotation of the DSMZ type-strain, *Segatella copri* DSM 18205, the essential enzymes for the tricarboxylic acid cycle (TCA), alpha-ketoglutarate dehydrogenase, succinyl-CoA synthetase and succinate dehydrogenase were not identified, despite the remaining genes involved in the TCA pathway being present. Absent genes and catalytic function were shaded with a *.

### Proposal of novel subspecies of *eno*(*-*) *Segatella copri*

A further five *S. copri* strains were found not to contain complete glycolytic and gluconeogenic pathways via Prokka genome annotation, including the *S. copri* DSM 18205 duplicate *S. copri* JCM 13468. The strains without an identified *eno* are three DSMZ non-type strains, *S. copri* DSM 18205, *S. copri* DSM 108494 (LKV-178-WT-2C), *S. copri* DSM 111828 (RHA03), and two NCBI available strains, *S. copri* RHA01 and *S. copri* RHA02 (Supplementary Sheet 1). *S. copri* DSM 108494 was isolated from 3-month old pig faeces, and *S. copri* DSM 18205, *S. copri* DSM 111828, *S. copri* RHA01 and *S. copri* RHA02 were isolated from adult human faeces.

A whole genome phylogenetic tree of all the collected *S. copri* isolates (*n* = 42) was produced and revealed that the *S. copri* strains lacking *eno* (DSM 18205, JCM 13468, LKV-178-WT-2C, RHA03, RHA01 and RHA02) were clustered into a subspecies (Figure 2). *S. copri* N-01 was grouped into the subspecies but contained two different enolases; enolase – *eno*, and enolase-phosphatase E1 (Figure 2).

**Figure 2:**
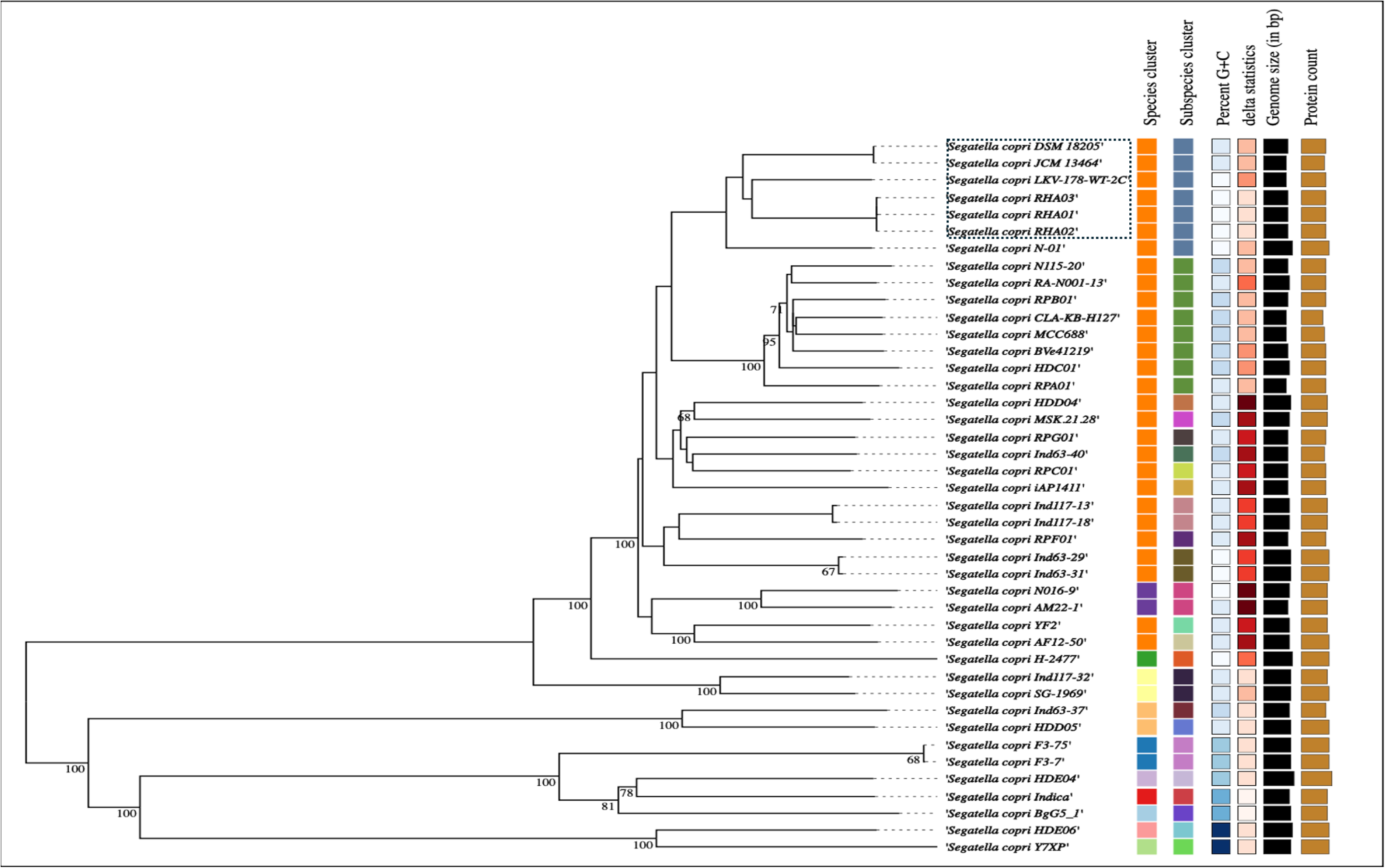
The whole genome phylogenetic tree of the NCBI accessed *Segatella copri* strains aligned with the DSMZ type-strain *Segatella copri* DSM 18205 has categorised the forty two *S. copri* isolates into one of eleven species clusters. *S. copri* isolates were further organised into one of twenty five subspecies clusters. The *eno*(-) *S. copri* isolates are highlighted with a dashed box; *S. copri* DSM 18205, JCM 13464, LKV-178-WT-2C, RHA03, RHA01 and RHA02.

The consistent absence of the key TCA cycle enzymes, alpha-ketoglutarate dehydrogenase, succinyl-CoA synthetase and succinate dehydrogenase across *S. copri* isolates may suggest a less recent evolutionary history than the loss of *eno* (Figure 2).

The absence of key enzymes involved in glycolysis, gluconeogenesis and TCA across *S. copri* could be due to the presence and increased activity of a metabolic pathway with the capacity to produce PEP, such as *ppdK* or *pckA* (Figure 2) (Koendjbiharie et al. 2021). The idea of *ppdK* or *pckA* being more active in the subspecies of *S. copri* may be due to these isolates being within environments with relative levels of compounds they are capable of converting to pyruvate or oxaloacetate and then PEP (Koendjbiharie et al. 2021). Hypothetically, the absence of TCA enzymes could have occurred and not impacted the cell’s fitness due to the presence of an alternative pathway to produce oxaloacetate, such as *pckA* producing oxaloacetate from malate (Koendjbiharie et al. 2021). And the loss of *eno* may have occurred due to the cell not being limited in its capacity to produce PEP, via both *pckA* and *ppdK* simultaneously (Koendjbiharie et al. 2021).

The alternative microbial synthesis of PEP intestinally via both *pckA* and *ppdK* has been predominantly associated with microbes surviving nutrient limited environments via regulating their carbon flux through non-carbohydrate activated gluconeogenesis (Rhode et al. 2012; Basu et al. 2018; Sommer and Newell 2019). The capacity of *S. copri* to regulate its carbon flux via non-carbohydrate activated gluconeogenesis would have host related health outcomes ranging from metabolic benefit to pathogen proliferation (Long et al. 2017; Bertin et al. 2014; Rhode et al. 2012; Basu et al. 2018; Sommer and Newell 2019). These host health outcomes are comparable to the host benefits and issues associated with *S. copri* intestinal relative abundance, as *S. copri* has been correlated with glucose homeostasis in various *in vivo* models (Yang et al. 2024; Péan et al. 2020; Kovatcheva-Datchary et al. 2015; Liu et al. 2023; De Vadder et al. 2016; Basu et al. 2018; Sommer and Newell 2019). But also, the impact of non-carbohydrate activated gluconeogenesis upon microbial proliferation could be one of the routes in which associates *S. copri* with various intestinal inflammatory disorders (Long et al. 2017; Bertin et al. 2014; Rhode et al. 2012; Basu et al. 2018).

The host related outcomes of *S. copri*, in relative abundance, actively maintaining intestinal glucose levels via non-carbohydrate activated gluconeogenesis has not been shown and would require clear identification of the metabolic output and capacity of the surrounding intestinal microbiome. For example, the capacity of *S. copri* to increase intestinal glucose via *pckA* or *ppdK* PEP synthesis, would likely require the gluconeogenic function of another enolase containing microbes, as this work has identified the absence of enolase within *S. copri* as potential evolutionary factor (Table 1) (Koendjbiharie et al. 2021). The capacity of a microbe, such as *S. copri,* to synthesise PEP without enolase would provide the intestinal microbiome glucose from non-carbohydrate sources but will also offer the opportunity for microbes to engage in cross-feeding with *S. copri* (Koendjbiharie et al. 2021).

The increased transcription of *ppsA* has been linked with various intestinal disorders, predominantly associated with inflammation and haemorrhaging, however *ppsA* was not identified within any of the screened *S. copri* isolates (Long et al. 2017; Bertin et al. 2014). The identification of *ppsA* would create issues with utilising *S. copri* isolates as probiotics, in particular within the maternal system (Long et al. 2017; Bertin et al. 2014). The utilisation of *S. copri* isolates in the future must screen for *ppsA,* as its presence within *S. copri* would provide a similar metabolic benefit as *pckA* and *ppdK*, that may lead to *ppsA* being transcribed by *S. copri* if received via horizontal gene transfer and becoming a functional gene (Long et al. 2017; Bertin et al. 2014).

### Alternative phosphoenolpyruvate synthesis pathways in *Segatella copri* DSM 18205

The analysis of the queried metabolic pathways within the NCBI reference genome, *S. copri* DSM 18205, produced via gapseq, revealed the presence of enzymes that are involved in the alternative phosphoenolpyruvate synthesis pathways; which utilise formate, ferredoxin and fumarate to produce pyruvate, fumarate to produce malate, convert pyruvate or malate to oxaloacetate, and the subsequent conversions of pyruvate to phosphoenolpyruvate, and oxaloacetate to phosphoenolpyruvate (Table 3) (Koendjbiharie et al. 2021; Koendjbiharie et al. 2018; Zimmermann et al. 2021; Maleki et al. 2017; Zhang et al. 2024; Crain et al. 2014). From the Prokka annotation report, four alternative PEP synthesis pathways that utilise *ppdK* or *pckA* to synthesise PEP, from pyruvate or oxaloacetate, were identified in *S. copri* DSM 18205, that do not require the function of *eno*, alpha-ketoglutarate dehydrogenase, succinyl-CoA synthetase and succinate dehydrogenase, to yield pyruvate, oxaloacetate or phosphoenolpyruvate (Figure 1; Tables 1-3).

**Table 3:**
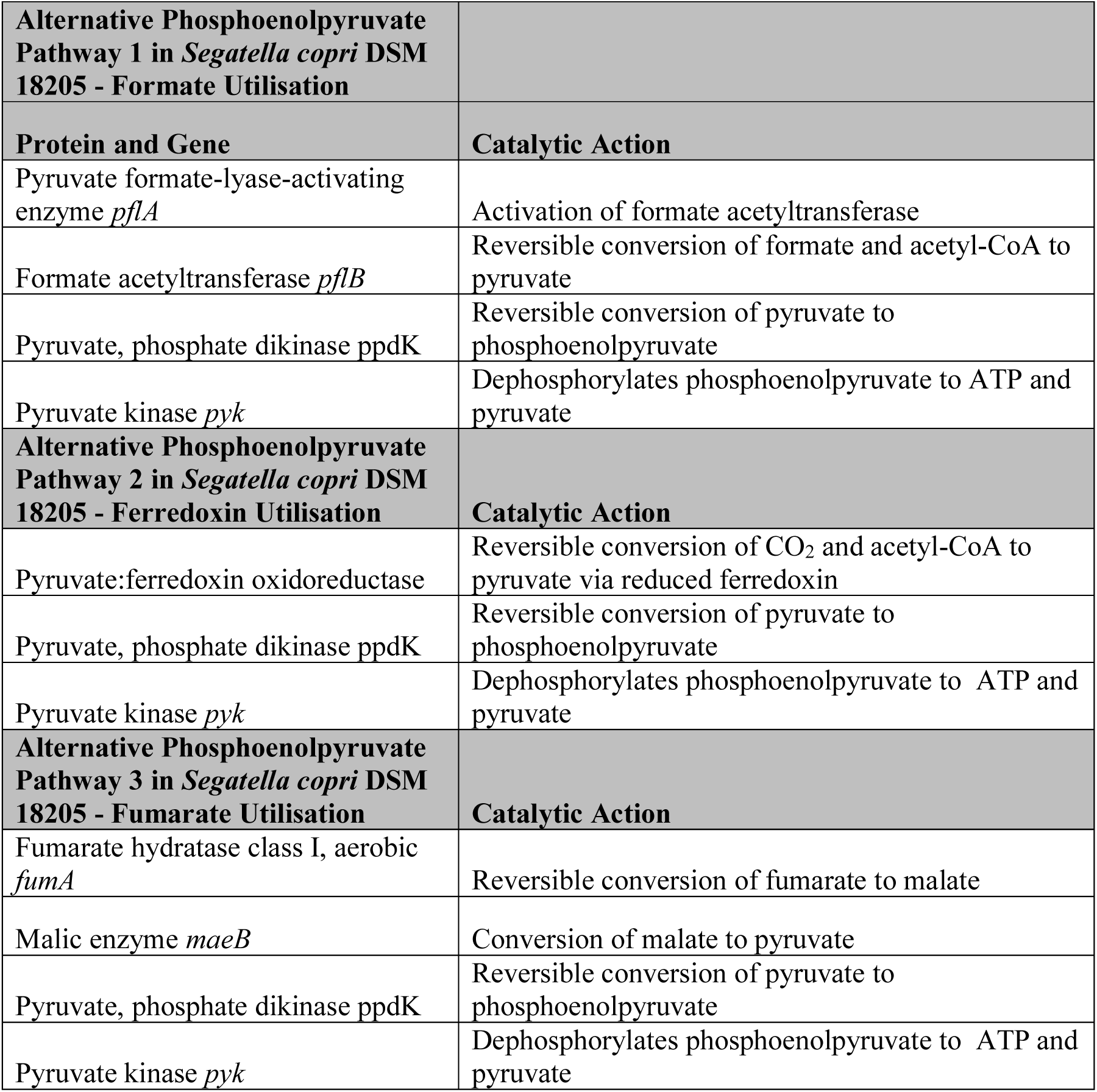
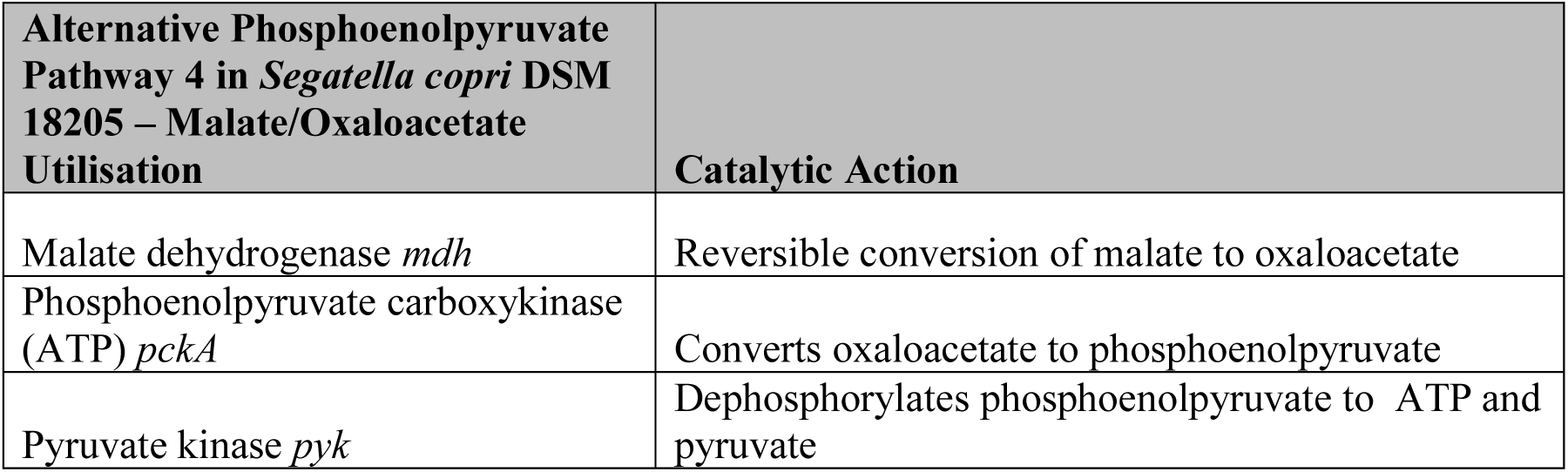
Through analysis of the Prokka genome annotation of the DSMZ type-strain, *Segatella copri* DSM 18205, four alternative phosphoenolpyruvate (PEP) pathways, that do not require enolase (*eno*), have been identified. The alternative PEP biosynthesis pathways bypass *eno* requirement via utilising one of two enzymes, Pyruvate, phosphate dikinase (*ppdK*) or Phosphoenolpyruvate (*pckA*), that can yield PEP from pyruvate and oxaloacetate, respectively.

The first alternative PEP synthesis pathway identified in *S. copri* DSM 18205 could function through the *pflAB* complex, with *pflA*, the pyruvate formate-lyase 1-activating enzyme, and *pflB,* the pyruvate formate-lyase enzyme (also known as formate C-acetyltransferase) (Figure 1; Table 3) (Maleki et al. 2017; Zhang et al. 2024; Crain et al. 2014; Seemann 2014). Pyruvate formate-lyase enzyme is considered crucial for anaerobic metabolism due to converting pyruvate coenzyme A into formate and acetyl-CoA (Maleki et al. 2017; Zhang et al. 2024; Crain et al. 2014). However, under anaerobic conditions with high formate, the reaction can also proceed in reverse and produce pyruvate from formate (Figure 1; Table 3) (Maleki et al. 2017; Zhang et al. 2024; Crain et al. 2014; Broderick et al. 2000; Caceres et al. 2023). Various *in vitro* studies on the metabolic activity of *S. copri* have found the production and export of formate is common in *S. copri* isolates and believed to occur during the production of acetyl-CoA from pyruvate by *pflB* (Franke and Deppenmeier 2018; Huang et al. 2021). However, within anaerobic environments, like the intestines, the *pflB* reaction occurs in the formate to pyruvate direction and indicates that with increased levels of formate within an intestinal environment, the potential for *pflAB* containing *S. copri* to produce pyruvate from formate is perceivable (Maleki et al. 2017; Zhang et al. 2024; Crain et al. 2014; Broderick et al. 2000; Caceres et al. 2023). The production of pyruvate from formate, although not the predominant direction of the *pflAB* reaction, can provide benefit to microbes by merely providing pyruvate for the conversion to formate and acetyl-CoA, or the conversion of pyruvate to PEP by *ppdK* (Figure 1; Table 3) (Maleki et al. 2017; Zhang et al. 2024; Crain et al. 2014; Broderick et al. 2000; Caceres et al. 2023; Gupta et al. 2021; Melkonian et al. 2019).

While the *pflB* reaction may be dependent on environmental factors, such as oxygen and substrate level, the activation of *pflB* is known to require SAMe and a reduced flavodoxin (Bonitatibus et al. 2025; Zhang et al. 2024; Troitzsch et al. 2024; Boswinkle et al. 2022). The genes *metK* and *fldA*, involved in the synthesis of SAMe and flavodoxin, respectively, were identified within *S. copri* DSM 18205 genome, and may indicate the capacity to synthesise these compounds during the activation of *pflB* (Bonitatibus et al. 2025; Zhang et al. 2024; Troitzsch et al. 2024; Boswinkle et al. 2022). Although the reduction of flavodoxin via PFOR requires TPP and the capacity of *S. copri* to produce TPP requires further investigation, as the activation of PFOR by *S. copri* without the ability to endogenously produce TPP could incite the consumption of intestinal TPP by *S. copri* (Bonitatibus et al. 2025; Zhang et al. 2024; Katsyv et al. 2021).

The second alternative PEP synthesis pathway identified in *S. copri* DSM 18205 would also access the PPO node via pyruvate synthesis but requires the direct action of PFOR and would be directly impacted by TPP (Katsyv et al. 2021). This pathway may occur through the reversed action of PFOR, with PFOR found to operate in both directions in energy requiring microbes (Figure 1; Table 3) (Bonitatibus et al. 2025; Furdui and Ragsdale 2000; Dietrich and Muller et al. 2023). It was reported that PFOR could utilise ferredoxin as an electron carrier during the conversion of acetyl-CoA and CO_2_ to pyruvate during microbial energy synthesis, providing an alternative avenue for pyruvate synthesis for PFOR and containing microbes (Figure 1; Table 3) (Bonitatibus et al. 2025; Furdui and Ragsdale 2000; Dietrich and Muller et al. 2023). Minimal work has investigated the capacity of microbes, apart from some acetogens, to utilise PFOR bidirectionally, although the reverse function of PFOR in *ppdK* containing microbes may provide additional pyruvate for PEP synthesis (Bonitatibus et al. 2025; Furdui and Ragsdale 2000; Dietrich and Muller et al. 2023). It is also possible for a *pflB* and PFOR containing microbe to utilise the acetyl-CoA from the *pflB* conversion of pyruvate to formate, which may link these reactions in a pyruvate and acetyl-CoA exchange, but these reactions may also directly impact one another due to both requiring the reductive action of PFOR and TPP as a cofactor (Zhang et al. 2024; Bonitatibus et al. 2025; Furdui and Ragsdale 2000; Dietrich and Muller et al. 2023).

The third and fourth alternative PEP synthesis pathways identified in *S. copri* DSM 18205 could occur via the conversion of fumarate to malate by *fumA*, with malate being converted to either pyruvate or oxaloacetate by *maeB* and *mdh,* respectively (Koendjbiharie et al. 2021; Wei et al. 2023). The production of pyruvate or oxaloacetate from malate would allow *S. copri* DSM 18205 to enter the PPO via either *ppdK,* if the malate is converted to pyruvate, or *pckA* if the malate is converted to oxaloacetate (Koendjbiharie et al. 2021) (Figure 1).

Within *S. copri* DSM 18205, a pyruvate kinase (*pyk*) was observed, indicating the capacity to dephosphorylate PEP produced by both *ppdK* and *pckA* reactions (Figure 1; Table 3) (Gupta et al. 2021; Melkonian et al. 2019; Chiba et al. 2015). During the production of ATP via the dephosphorylation of PEP, the by-products pyruvate and Pi are produced (Figure 1) (Koendjbiharie et al. 2021; Chiba et al. 2015). The production of ATP and pyruvate also provides the opportunity for the PPO node to be accessed via *ppdK* (Figure 1) (Koendjbiharie et al. 2021; Cline et al. 2011; Llamas-Ramirez et al. 2020; Koendjbiharie et al. 2018; Saur et al. 2005).

The requisition of phosphates by *ppdK* and *pckA* during PEP synthesis has been observed to be impacted by the presence of Pi, with both *ppdK* and *pckA* found to receive phosphates from Pi or GTP over ATP in the presence of Pi (Figure 1) (Chastain et al. 2011; Tjaden et al. 2006). The ability of the *ppdK* and *pckA* to not consume ATP during PEP synthesis may create a substrate cycle with an ATP yield and could be a key metabolic pathway within *S. copri* DSM 18205 due to the presence of *ppdK* and *pckA* (Figure 1) (Koendjbiharie et al. 2021; Chastain et al. 2011; Tjaden et al. 2006).

### Access to the phosphoenolpyruvate-pyruvate-oxaloacetate node with an ATP Yield in *Segatella copri* DSM 18205 genome

Likened to other substrate cycles, the PPO node commonly amounts in no net yield of pyruvate, PEP or ATP (Koendjbiharie et al. 2021; Chiba et al. 2015). During the possible PPO substrate cycle within *S. copri* DSM 18205, pyruvate and oxaloacetate is converted directly to PEP via *ppdK* and *pckA,* respectively, in an ATP requiring reactions, then PEP is dephosphorylated to pyruvate, Pi and ATP by pyruvate kinase, in turn providing the pyruvate and ATP to reproduce PEP via *ppdK* (Figure 1) (Koendjbiharie et al. 2021; Chiba et al. 2015). However, due to the production of a Pi during the dephosphorylation of PEP, the production of PEP by *ppdK* and *pckA* may not require ATP, as *ppdK* and *pckA* have been observed to utilise Pi and GTP instead of ATP in the presence of Pi (Figure 1) (Koendjbiharie et al. 2021; Chiba et al. 2015; Chastain et al. 2011; Tjaden et al. 2006). The PPO cycle may provide increased fitness to microbes such as *S. copri*, with the capacity to utilise end products as substrates to produce PEP, whilst yielding ATP (Figure 1) (Koendjbiharie et al. 2021; Chiba et al. 2015; Chastain et al. 2011; Tjaden et al. 2006). Albeit the production and dephosphorylation of PEP via this pathway may not be the only metabolic benefit received, as PEP may be alternatively shuffled into other pathways, while the PEP being utilised in gluconeogenesis would be improbable due to the lack of an *eno* in *S. copri* DSM 18205 (Table 1) (Koendjbiharie et al. 2021; Zhang et al. 2024; Gupta et al. 2021; Melkonian et al. 2019; Chiba et al. 2015; Chastain et al. 2011; Tjaden et al. 2006).

The additional production of PEP in bacteria is often associated with the phosphotransferase system, which allows bacteria to utilise PEP as a phosphate donor to transfer sugars into the cell with increased efficiency (Siebold et al. 2001; Deutscher et al. 2014) However, PEP:PTS interaction requires a complement of enzymes, with specific functions in phosphorylating and transferring sugars into the cell, including Enzyme I, Histidine phosphocarrier protein, Enzyme IIA, Enzyme IIB and Enzyme IIC, which was not identified in the *S. copri* DSM 18205 genome during the analysis of the Prokka annotation (Deutscher et al. 2014; Patel et al. 2006; Deutscher et al. 2006). Although, a galactinol-specific Enzyme IIC (*gatC*) was not identified within the *S. copri* DSM 18205 genome, however without the entire complement of PTS enzymes, *S. copri* DSM 18205 does not have the genomic capacity to utilise PEP via the PTS system and suggests an alternative fate of additional PEP produced by *S. copri* DSM 18205 (Koendjbiharie et al. 2021; Gupta et al. 2021; Melkonian et al. 2019; Deutscher et al. 2014; Patel et al. 2006; Deutscher et al. 2006).

Although the action of a PPO substrate cycle in *S. copri* DSM 18205 cannot be confirmed bioinformatically, the genomic capacity to utilise formate, ferredoxin and fumarate to synthesise pyruvate and oxaloacetate, and the conversion of pyruvate and oxaloacetate to PEP was determined in the NCBI reference strain, *S. copri* DSM 18205 (Table 3). The subsequent production of pyruvate, Pi and ATP during PEP dephosphorylation, was also determined and may further increase PPO node activity (Table 3) (Koendjbiharie et al. 2021; Zhang et al. 2024).

### Varied rates of conservation of alternative phosphoenolpyruvate synthesis pathways across *Segatella copri* subspecies

To establish if the genes involved in accessing the PPO node via formate, ferredoxin and fumarate utilisation are conserved across the NCBI access *S. copri* isolates, the level of relatedness of the selected isolates was determined through a whole genome based phylogenetic tree (Figure 2) (Van Damme et al. 2022; Martin et al. 2003; Luo et al. 2015). Whilst the evolutionary network and alignment data indicates that various of the selected *S. copri* isolates are not the same species as *S. copri* DSM 18205, the alignment of gene sequences of divergent species may indicate a level of genetic function through the conversation of sequences after evolutionary divergence (Figure 2) (Van Damme et al. 2022; Baum et al. 2008; Maharjan et al. 2012; Koonin et al. 2003; Parsch et al. 2005; Frazer et al. 2003).

To demonstrate the conservation of enzymes involved in a potential PPO substrate cycle via formate, ferredoxin and fumarate utilisation within the *S. copri* isolates, the nucleotide sequences of the genes encoding the enzymes *metK, fldA,* PFOR, *pflA, pflB, fumA, maeB, ppdK, mdh, pckA* and *pyk,* from the isolates were assessed for homology with the respective gene sequences in *S. copri* DSM 18205 (Table 3; Figure 3) (Koendjbiharie et al. 2021; Gupta et al. 2021; Melkonian et al. 2019; Zhang et al. 2024; Van Damme et al. 2022; Baum et al. 2008; Maharjan et al. 2012; Koonin et al. 2003; Parsch et al. 2005; Frazer et al. 2003). The sequence homology heatmap was aligned with the phylogenetic tree of the *S. copri* isolates, to allow for the potential negative trend of gene sequence conservation and evolutionary divergence to be viewed (Table 3; Figure 3) (Koendjbiharie et al. 2021; Gupta et al. 2021; Melkonian et al. 2019; Zhang et al. 2024; Van Damme et al. 2022; Baum et al. 2008; Maharjan et al. 2012; Koonin et al. 2003; Parsch et al. 2005; Frazer et al. 2003; Omer et al. 2017; Lato et al. 2021).

**Figure 3:**
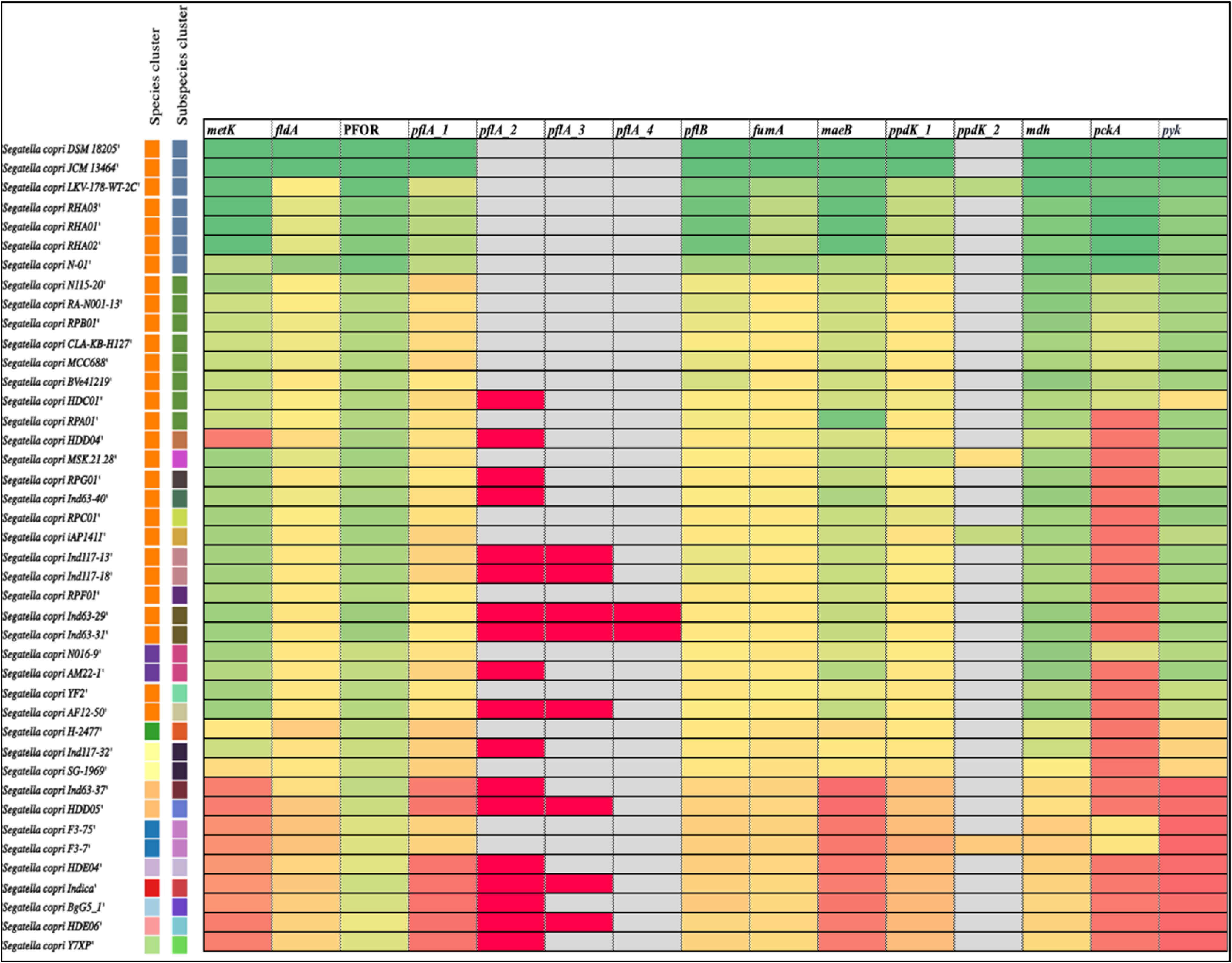
The whole genome phylogenetic tree ordered *Segatella copri* strains, in species and subspecies clusters, were aligned with a heatmap produced from the percentage identity of the genes conferring to the enzymes involved in the alternative phosphoenolpyruvate (PEP) synthesis pathways via the utilisation of formate, ferredoxin and fumarate.

The highest level of conservation for all alternative PEP synthesis pathways was seen in the *S. copri* subspecies which lacked *eno* (Figure 2; Figure 3). Whilst speciation is not possible from phylogenetics alone, the potential for these alternative PEP synthesis pathways to be viewed as factors in the evolution of this subspecies is possible (Figure 3). The most conserved gene across all subspecies of *S. copri* was PFOR and may indicate an interaction between *Segatella* spp. and ferredoxin is present, but also that all isolates require TPP to utilise PFOR (Figure 3). A positive relationship between *S. copri* and ferredoxin could explain the positive effects of plant based products upon the relative abundance of *S. copri,* as ferredoxins are highly present within chlorophyll, while *Segatella* spp. have been suggested to have the capacity to produce TPP (Blanco-Miguez et al. 2023; Tournaire et al. 2023). Although this work did not do look into the genomic capacity of the *S. copri* isolates to produce TPP, the high level of conservation of PFOR across the isolates indicates that TPP synthesis by *S. copri* must be investigated and could detail if TPP synthesis is associated with PFOR function (Katsyv et al. 2021).

The *pflA* gene appears to have undergone genetic duplication in isolates (RPG01, HDD04, HDC01, Ind63-40, HDE06, BgG5_1, Y7XP, AM22-1, Ind63-37, Ind117-32 and HDE04), with multiple duplications observed in (Ind63-31, AF12-50, Ind117-18, Ind117-13, Ind63-29, Indica and HDD05) (Figure 3). Although the level of conservation is not significant for the duplicated genes and may have an alternative function (Figure 3). Additional potential duplication events were observed in the isolates iAP1411, MSK.21.28, LKV-178-WT-2C and F3-7, which were identified with two copies of *ppdK,* both with high levels of homology to the *S. copri* DSM 18205 *ppdK* sequence and may suggest an increased capacity to produce PEP from pyruvate within these isolates (Figure 3) (Koendjbiharie et al. 2021; Tian et al. 2003; Pearson et al. 2013). The intra-species sequence retention of *pflA* and *ppdK* suggests a potential dependency on these genes due to high sequence conservation despite the presence of multiple copies of the gene and evolutionary divergence (Figure 3) (Koendjbiharie et al. 2021; Zhang et al. 2024; Parsch et al. 2005; Frazer et al. 2003; Omer et al. 2017; Tian et al. 2003; Pearson et al. 2013).

Further work is required to understand the central metabolism of the subspecies of *S. copri* which lack key central metabolism genes. The transcriptomic and metabolomic analysis of *S. copri* isolates in formate, ferredoxin or fumarate rich environments could reveal the relationship between these electron carrying compounds and *S. copri.* The understanding of this access to the PPO node by *S. copri* via various sources may further detail the metabolic potential of intestinal bacteria.

### Increased conservation of protein sequences indicates conserved function in *Segatella copri* isolates

Although nucleotide sequence conservation can suggest function, the functionality of a gene is dictated by the protein produced. The alignment of the respective proteins transcribed from genes was analysed for further insight into the potential PEP synthesis pathway via formate, ferredoxin and fumarate in the *S. copri* isolates (Joshi et al. 2007; Koonin et al. 2002; Jensen et al. 2003). The protein sequence alignment indicated that the protein sequences produced from the highly conserved nucleotide sequences for the genes, *metK, fldA,* PFOR, *pflA, pflB, fumA, maeB, ppdK, mdh, pckA* and *pyk,* were even more conserved than the nucleotide sequences, indicating not only a similar function of these respective proteins but also that these proteins are conserved in these divergent isolates potentially due to an evolutionary fitness (Jensen et al. 2003; Siltberg-Liberles et al. 2011; Huang et al. 2013).

The respective protein sequences of all the *pflA* duplicates that were found not significantly similar via nucleotide alignment, were observed to share >25% sequence homology with the *S. copri* DSM 18205 *pflA* protein sequence. For the isolates, that were found with duplicate *pflA* sequences, it was observed that one of two of the duplicate sequences was highly conserved at >95% PI, while the other duplicate *pflA* was less similar at ∼25% PI, and could indicate an alternative function or lack of function in the dissimilar *pflA* duplicates (Wilson et al. 2000; Chung et al. 1996; Rost et al. 1999). The protein alignment also revealed the functional integrity of the duplicate *ppdK* in isolates (iAP1411, MSK.21.28, LKV-178-WT-2C and F3-7), potentially giving these isolates two functional *ppdK* proteins with an increased capacity to synthesis PEP from pyruvate and dephosphorylate PEP.

### Concluding Remarks

Past work has investigated the growth of *S. copri* DSM 18205 on glucose and demonstrated that glucose was consumed during the *in vitro* growth experiment utilising peptone yeast glucose (PYG) broth (Franke and Deppenmeier 2018; Huang et al. 2021). The same work also sequenced *S. copri* DSM 18205 and reported the absence of *eno* during genome annotation (Franke and Deppenmeier 2018). The capacity of microbes to yield energy from glucose without completing glycolysis is not clear. Although the interchange of 2-phosphoglycerate by glycolytic and gluconeogenic pathways, may allow for partial cycling of these pathways, although the lack of *eno* indicates PEP will not be a product produced (Table 1) (Chaudhry et al. 2018). Despite PEP not being produced from this possible 2-phosphoglycerate interaction of glycolytic and gluconeogenic pathways, the interchanging between these two incomplete pathways may still have the capacity to yield 4 ATP per cycle, with ATP yielded by the phosphoglycerate kinase (*pgk*) reaction in glycolysis and gluconeogenesis (Table 1) (Chaudhry et al. 2018).

Whilst the observations are based on genomic and proteomic analysis, prior research has observed that *S. copri* relative abundance increased significantly in media with increased formate during *in vitro* experiments, though the study did not specifically quantify the extent of increased growth with higher levels of formate (Parkar et al. 2021). Research has not looked directly at the impacts of formate on *S. copri* growth, but various studies have highlighted the inverse intestinal relationship of *Segatella* spp. with bacteria that have the capacity to utilise formate at high rates during acetogenesis and methanogenesis (Kumpitsch et al. 2021; Sun et al. 2021; Karakashev et al. 2006; Prochazkova et al. 2023; Aryee et al. 2023; Aguillar et al. 2020; Trischler et al. 2022; Laverde Gomez et al. 2019; Wang et al. 2023).

The utilisation of fumarate as an electron acceptor and production of fumarate as an intermediate product have been reported in *Segatella* spp. (Franke and Deppenmeier 2018). While the capacity to produce fumarate during TCA is not likely in *S. copri* DSM 18205 due to the lack of succinate dehydrogenase within the genome (Table 2). The consumption of fumarate is often attested to acetogenic and methanogenic microbes and could be also contributing to the negative relationship between *Segatella* spp. and acetogens and methanogens (Asanuma et al. 1999; Aryee et al. 2023; Aguillar et al. 2020).

The utilisation of ferredoxin as an electron carrier by PFOR predominantly occurs during the synthesis of acetyl-CoA but has been shown to be bidirectional and synthesise pyruvate (Bonitatibus et al. 2025; Furdui and Ragsdale 2000; Dietrich and Muller 2023). The actions of PFOR in *S. copri* have been previously explained but have only focused on the capacity of PFOR to utilise ferredoxin to produce acetyl-CoA (Franke and Deppenmeier 2018). Although indirectly correlated, the negative association of *Segatella* spp. and methane levels, can suggest with knowledge of the conserved PPO cycle within the screened *S. copri* isolates, that the negative association observed between *Segatella* spp. and methane could potentially be due to competition with the methanogens and acetogens (Koendjbiharie et al. 2021; Zhang et al. 2024; Tian et al. 2003; Pearson et al. 2013; Kumpitsch et al. 2021; Sun et al. 2021; Karakashev et al. 2006; Prochazkova et al. 2023; Aryee et al. 2023; Aguillar et al. 2020; Trischler et al. 2022; Laverde Gomez et al. 2019; Wang et al. 2023; Parkar et al. 2021).

Whilst this work has conceptualised the idea that *Segatella* spp. may receive metabolic benefit from the utilisation of electron donors via biochemical pathways that are not likened to methanogenic or acetogenic activity. It has also raised the prospect that the product of electron donor utilisation within intestinal environments could range from being consumed during acetogenesis and methanogenesis, and leading to the production of methane, or alternatively be utilised by *Segatella* spp. to synthesise PEP and PEP-intermediates.

Although methane emissions from ruminants have been identified as a factor contributing to global greenhouse gas emissions, the increased activity of methanogens and acetogens within the human intestinal microbiome is also known to increase the production of methane and is linked to constipation-predominant Irritable Bowel Syndrome (IBS-C) (Wagar and Rehan 2019; Goshai et al. 2016). Alternatively, within intestinal environments with a low relative abundance of electron donor consuming microbes, the accumulation of electron donors intestinally is known to cause gastrointestinal issues from dysbiosis, higher intestinal permeability and restricted host metabolism (Winter and Bäumer, 2023; Kwon et al. 2016). However, the consumption of electron donors by acetogenic microbes does have the capacity to positively impact the host and is dependent upon the microbes cross-feeding with acetogens, as the acetate produced by acetogens can be utilised by various gut bacteria to synthesise butyrate, oppose to the acetate being converted to methane (Laverde Gomez et al. 2018; Trischler and Müller 2026; Daisley et al. 2021).

*In vitro* growth analysis with transcriptomic and metabolomic analysis could clarify whether the presence of formate, ferredoxin and fumarate influence PEP synthesis in *S. copri* (Franke and Deppenmeier 2018; Huang et al. 2021). Analysing pyruvate, oxaloacetate and phosphoenolpyruvate levels, in partisan with formate, ferredoxin and fumarate levels will provide insight into the interactions between *S. copri* with these compounds (Huang et al. 2021; Polyak et al. 2003; Higgs et al. 2001; Varik et al. 2017). The ability of *S. copri* to yield PEP and ATP from formate, ferredoxin or fumarate would provide insight into new prebiotics to increasing *S. copri* intestinally, which may also be beneficial for other probiotic microbes (Koendjbiharie et al. 2021; Zhang et al. 2024; Parkar et al. 2021). While formate, ferredoxin and fumarate are regularly present within the human intestine, the benefit of limiting the colonising capacity of competing microbes, such as acetogens and methanogens, may also provide the opportunity for these compounds to be converted to pyruvate or oxaloacetate, and utilised to produce PEP by other microbes (Koendjbiharie et al. 2021; Zhang et al. 2024; Kumpitsch et al. 2021; Sun et al. 2021; Prochazkova et al. 2023; Aguillar et al. 2020; Trischler et al. 2022; Parkar et al. 2021). Alternatively, understanding the relationship between *S. copri*, these electron carrying molecules, and acetogenic and methanogenic bacteria, must be investigated to the reveal if *S. copri* growth is negatively impacted by the consumption of these compounds by acetogenic and methanogenic bacteria intestinally (Koendjbiharie et al. 2021; Zhang et al. 2024; Kumpitsch et al. 2021; Sun et al. 2021; Prochazkova et al. 2023; Aguillar et al. 2020; Trischler et al. 2022; Parkar et al. 2021). The *in vitro* analysis of *S. copri* strains capacity to utilise formate, ferredoxin and fumarate must consider the central metabolism subspeciation of *S. copri* isolates, with the expectation to observe varied metabolic capacities across the subspecies of *eno*(*+*) and *eno*(-) *S. copri*.

The likely reduced capacity to utilise monomers by the *eno*(-) *S. copri* isolates may lead to an increased susceptibility to competition within the intestines (Koendjbiharie et al. 2021; Chaudhry et al. 2018). The loss of *eno* may be an evolutionary response to the intestinal environment the *eno*(-) *S. copri* isolates reside in but could be also due to the cultural shifts we as a human society have undergone, with diet, hygiene and socio-economics all being factors seen to impact the intestinal microbiome (Shridhar et al. 2024; Conlon and Bird 2014). Alternatively, the retention of *eno* within the *eno*+ *S. copri* isolates could be due to the requirement of monomer utilisation within the intestines (Koendjbiharie et al. 2021; Chaudhry et al. 2018).

The impacts within the intestines of these alternative PEP synthesis pathways within *S. copri* may have already been identified, with the relative abundance of PEP and *S. copri* independently reported to mediate intestinal bacteria but also inflame intestinal epithelia (Nieman et al. 2025; Lo Presti et al. 2023; Chen et al. 2021; Yin et al. 2024). Moreover, the interaction of other constituents of the intestinal microbiome and the exported PEP may result in excess intestinal PEP being converted to glucose, if the microbes contain an enolase and gluconeogenic function (Koendjbiharie et al. 2021; Melkonian et al. 2019). Whilst it is unlikely for the *eno*(-) *S. copri* isolates to shuffle PEP into the gluconeogenic pathway, the capacity of gluconeogenic bacteria to yield glucose from intestinal PEP is feasible (Koendjbiharie et al. 2021; Melkonian et al. 2019). This ability to convert PEP yielded by *S. copri* from formate, ferredoxin or fumarate, into glucose by glucose requiring microbes may work to lower community wide glucose competition within the intestines (Koendjbiharie et al. 2021; Melkonian et al. 2019). The capacity of the intestinal microbiome to produce glucose from *S. copri* exported PEP may facilitate the intestinal glucose homeostasis, with glucose regulation reported to be associated with *S. copri* abundance in diabetic animal models (Yang et al. 2024; Pean et al. 2020). Furthermore, the observed increased fat accumulation in pigs with a higher relative abundance of *S. copri* could also be due to the increased conversion of exported PEP to glucose by other intestinal microbes, which could be absorbed by intestinal epithelia and converted to fat within host cells (Chen et al. 2021).

In conclusion, this work has identified the evolutionary subspeciation of the type-strain *S. copri* DSM 18205 and other *eno*(-) *S. copri* isolates (Figure 2). It was also seen that the evolutionary divergence of the *eno*(-) subspecies may be explained by the increased conservation of the alternative PEP pathways in the *eno*(-) subspecies *S. copri* (Table 3; Figure 2). While this work has established the genomic capacity of the *S. copri* isolates to produce pyruvate or oxaloacetate from formate, ferredoxin and fumarate, and subsequently access the PPO node via *ppdK* or *pckA*, the PFOR enzyme was the most conserved across all *S. copri* isolates, which is a reaction depending directly on thiamine pyrophosphate (TPP) availability (Table 3; Figure 3) (Katsyv et al. 2021). The catalytic requirement of TPP does suggest that if *S. copri* is producing utilising PFOR in either to reduce flavodoxin or ferredoxin, that the *S. copri* isolates would need to sequester intestinal TPP or have the capacity to synthesise its own TPP (Katsyv et al. 2021). Various studies have reported the synthesis of TPP via *Segatella* spp. and could be a metabolic output that fluctuates with the metabolic requirement of the microbe, i.e. flavodoxin or ferredoxin availability (Katsyv et al. 2021; Wan et al. 2022). Moreover, the reactions catalysed by *mdh* do not require TPP but are regulated by TPP and TPP derivatives binding to *mdh* (Mezhenska et al. 2020). Although TPP is not required for the catalytic functions of *pflA, pflB*, *fumA, maeB, ppdK, pckA* or *pyk,* TPP would be necessary for the increased growth and cellular proliferation that may occur in a microbe accessing the PPO node with an ATP yield (Koendjbiharie et al. 2021; Kaźmierczak-Barańska et al. 2025).

The investigation of TPP synthesis must occur in conjunction with the *in vitro* analysis of *S. copri,* and its capacity to access the PPO node via formate, ferredoxin and fumarate utilisation, as the absence of TPP synthesis within *S. copri* could indicate that *S. copri* abundance may lower intestinally TPP levels (Marrs and Lonsdale 2021). Alternatively, if *S. copri* is determined to be capable of synthesising TPP, the production and export of TPP intestinally by *S. copri*, would have intestinal community and host benefit through the acquirement of TPP, which is required for numerous metabolic reactions in all cells (Kaźmierczak-Barańska et al. 2025; Marrs and Lonsdale 2021).

## Abbreviations

(Pyr): Pyruvate,
(PEP): Phosphoenolpyruvate,
(ATP): Adenosine triphosphate,
(GTP): Guanosine triphosphate,
(Pi): Inorganic phosphate,
(PPO): PEP-Pyruvate-Oxaloacetate,
(TCA): tricarboxylic acid,
(PTS): Phosphotransferase system,
(WLP): Wood-Ljungdahl pathway,
(PEPCK): Phosphoenolpyruvate kinase,
(ANI): Average nucleotide identity,
(PI): Percentage identity,
(HPLC): High power liquid chromatography.

## Acknowledgements

Many thanks to researchers past and present who have contributed to the wealth of available genomic data online, with stringent metadata and methodology recorded. While writing down our history was a necessary cultural step in human evolution, the recording of bioinformatic data may be our next step.

Thanks to my supervisors and lab members at La Trobe University, that have allowed engaging conversations to evolve into rational arguments.

## Author CRediT Contributions

Luke Bosnar: Conceptualisation, data curation, formal analysis, investigation, methodology, visualisation, writing – original draft and writing – review and editing. Anya Shindler: Methodology, funding acquisition, project administration, resources, supervision and writing – review and editing. Ashley Franks: Methodology, funding acquisition, project administration, resources, supervision and writing – review and editing. Steve Petrovski: Supervision and writing – review and editing.

